# Familiarity with companions aids recovery from fear in zebrafish

**DOI:** 10.1101/098509

**Authors:** Ajay S. Mathuru, Annett Schirmer, Ng Phui Yeng Tabitha, Caroline Kibat, Ruey-Kuang Cheng, Suresh Jesuthasan

**Affiliations:** Yale-NUS College, Singapore; Institute of Molecular and Cell Biology, 61 Biopolis Drive, Singapore.; Department of Physiology, National University of Singapore, Singapore.; Department of Psychology, National University of Singapore, Singapore.; Neuroscience and Behavioral Disorders Program, Duke-NUS Graduate Medical School, Singapore.; Lee Kong Chian School of Medicine, Nanyang Technological University, Singapore

**Author notes:** Corresponding Author:*, *Mailing Address: 12 College Avenue West., #01-201, Singapore - 138610.

**Keywords:** Social buffering, fear, zebrafish, isotocin, cortisol

## Abstract

Interaction with social partners during or after a stressful episode aids recovery in humans and other mammals. We asked if a comparable phenomenon exists in zebrafish *(Danio rerio)* that live in shoals in the wild. In the first experiment, we observed that most quantifiable parameters of swimming behavior were similar when zerbafish swam alone or with companions. However, after exposure to an alarm substance (*Schreckstoff*), individuals recovering alone continued to display behaviors associated with fear after removal of the stimulus, while those recovering with companions did not. In the next two experiments, we examined the role of familiarity of companions. Subjects spent more time in the vicinity of familiar companions in a two-choice assay. While both familiar and unfamiliar companions reduced behavioral signs of distress, familiar companions additionally modulated cortisol and endogenous isotocin in subjects. Shortly after being united with familiar companions, isotocin spiked followed by a dampening of circulating cortisol levels. These results suggest that zebrafish experience fear attenuation in the presence of others and familiar companions are more effective at buffering the stress associated with escaping predation. Changes in behavior, circulating cortisol and isotocin levels due to social partners are reminiscent of changes due to amelioration of fear in some mammalian species in the presence of companions. The two phenomena may be related.

---

For many animals, living in groups is thought to be evolutionarily favorable compared to living alone ^1-3^. Although social interactions can be costly ^4^, individuals living in groups receive significant benefits including a lowered predation risk, efficient foraging, ease of finding mating partners, and support when caring for offspring ^3^. Furthermore, in mammals, close social contact with conspecifics has also been documented to reduce fear ^5^ (reviewed in ^6^) and to aid in recovery from stress ^6-8^. The phenomenon of improved recovery from aversive experiences in the presence of conspecifics has been termed “social buffering” ^6^. Positive social interactions are proposed to counter environmental stressors and the accompanying negative emotions in an individual via the release of opioids and oxytocin ^8-11^. However, our understanding of the molecular architecture, the neural circuits involved, and the evolutionary underpinnings of social buffering is still limited.

The dominant view to date is that the phenomenon of social buffering occurs in species that display social bonding in the form of mating pairs and/or care for offspring ^6,12^. This position rests on the fact that, at present, social buffering has been documented primarily in animals where these forms of social bonding are readily observed such as in some mammals and a few species of birds (^5,13,14^ and reviewed in ^6,7^). Though a single term has been used to describe social buffering in mammals, it can take more than one form. In squirrel monkeys for example, a mother can buffer infant stress ^15^ and an adult can reduce fear of the pair-bonded partner ^16^. In guinea pigs and humans on the other hand, social buffering properties are dependent on sex ^17,18^. In all cases however, the term implies active participation of partners when allowed. Still, it is unclear whether social buffering is unique to such species, in that it requires neurochemicals or brain structures that are absent in lower vertebrates. Moreover, if any form of social buffering is present in species that do not display strong social attachments, such as certain fish, either through passive or active participation of social partners, is still an open question.

In fishes, only a few studies have directly touched upon issues related to buffering ^19,20^. Although, the social nature of fish shoals is known and conspecific presence is considered important, increased cohesion in shoals to avoid predation has been interpreted primarily as exemplifying the “selfish-herd” effect ^21^. Authors of a study that reported recovery from immobility in a glucocorticoid receptor mutant zebrafish when a conspecific was visible, attributed the recovery to social buffering ^19^. However, whether such reports of buffering bear only a superficial similarity to the phenomenon previously described in mammals or reflects a conserved mechanism are only now beginning to be addressed ^22^.

Here we addressed this question by exploring buffering of fear in zebrafish, a promiscuous species that does not display parental care and is known to cannibalize newly fertilized eggs and larvae ^23^. We adapted the basic paradigm from research in mammals but assessed the subject’s recovery after a stressor or a predator threat was removed. We reasoned that this would allow us to dissociate social buffering from gregariousness directly due to a predator attack. Under such conditions, we find that distressed subjects shows signs of recovery from fear with companions that is absent when alone. We further reasoned that familiarity of companions should be of little consequence in such a scenario if only the “selfish herd” was operating, as any shoaling opportunity would reduce predation risk. Indeed, familiarity did not matter in attenuating fearful behavior of subjects. However, additionally, familiar companions appeared to be more effective in reducing the physiological measure of stress (whole-body circulating cortisol level). As anticipated by the role of oxytocin in amelioration of stress in mammals ^12,24,25^, we found increased central endogenous isotocin release (the fish homologue of oxytocin) when distressed zebrafish subjects were reunited with familiar conspecifics. Our results suggest that zebrafish experience attenuation of fear and the stress associated with detection of predators in the presence of familiar companions. The physiological processes mediating this buffering of fear are similar to those previously reported for rats, guinea pigs and humans.

## Results

### Behavioral markers of social buffering in the zebrafish

We explored the presence and properties of social buffering in zebrafish in four different experiments. For the first experiment, subjects were divided into four groups. Subjects in two groups were exposed to ordinary tank water (Control), while those in the other two groups were exposed to Schreckstoff ^26^, an alarm substance released from the skin of injured individuals that induces fear in others (Experimental). Subjects were then passed through a wash chamber and moved into a tank by themselves (Control-Alone and Experimental-Alone groups) or into a tank with four non-fearful conspecifics (Control-Conspecific and Experimental-Conspecific groups, Figure 1A). The behavior of subjects was recorded for 10 minutes and measures indicative of fear ^27^ were quantified using a custom-developed, semi-automated analysis method. Compared to subjects in the Control-Alone group, subjects in the Experimental-Alone group showed significantly more signs of fear (Figure 1B; Supplementary Video S1, S2). They spent more time in the bottom quarter of the tank (mean = 81.3%, 95 CI [70.8%, 91.8%], unpaired Welch’s t-test p < 0.001; Supp. Fig 1A), and showed more episodes of pausing (mean = 192.9, 95 CI [99.7, 286.0], unpaired Welch’s t-test, p = 0.01; Supp. Fig 1B), freezing (mean = 35.9, 95 CI [16.9, 54.9], unpaired Welch’s t-test p = 0.014; Supp. Fig 1C), and darting (mean = 2.3, 95 CI [3.6, 0.9], unpaired Welch’s t-test p < 0.007; Supp. Fig 1D).

**Figure 1.**
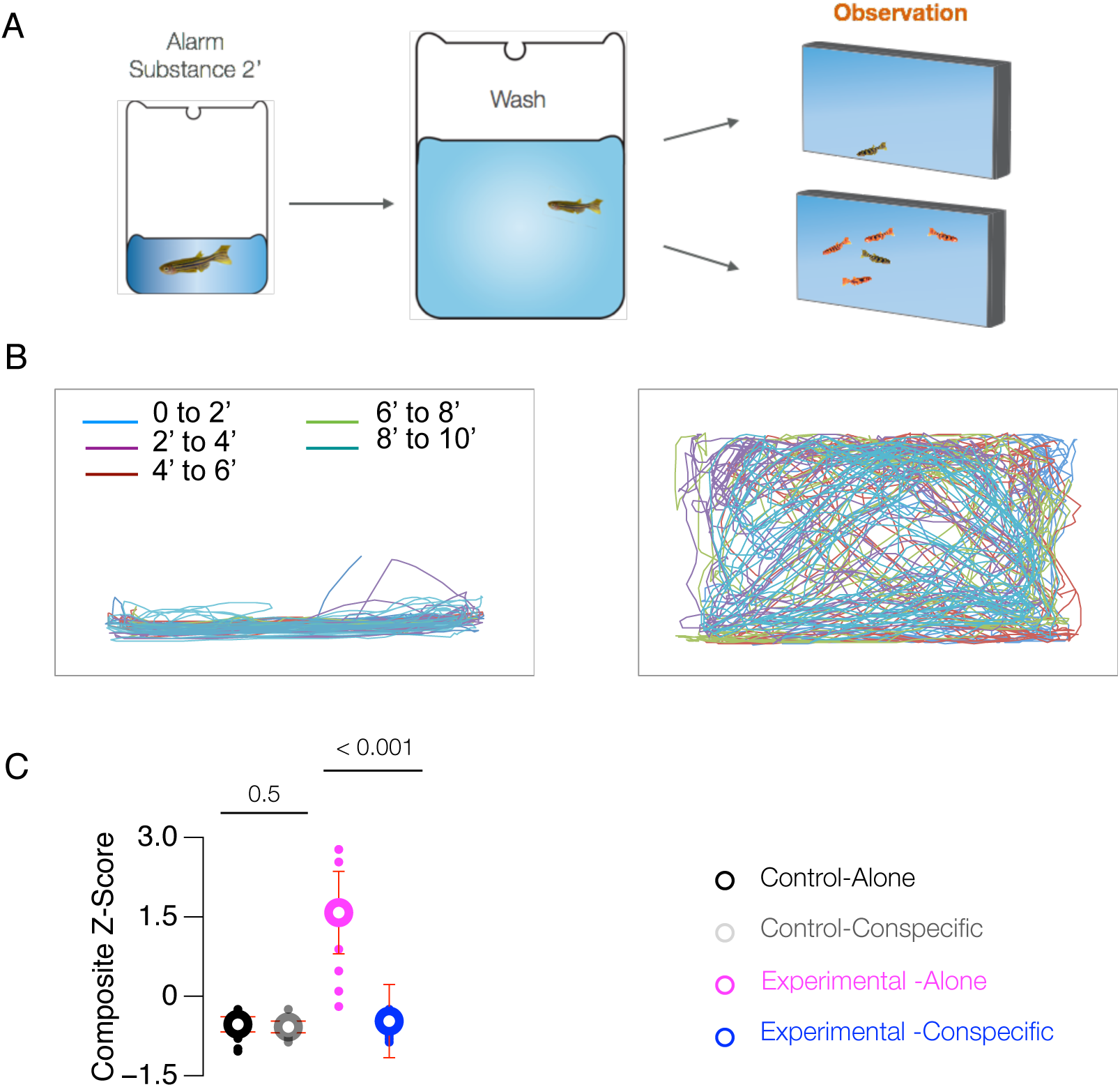
Behavioral markers of social buffering in the zebrafish. A) Schematic of social buffering paradigm. Experimental subjects were exposed to alarm substance for 2 minutes, washed and released into observation tanks for 10 minutes. B) Representative swimming traces (10 minutes) of Experimental subjects after exposure to alarm substance when alone (left) or with conspecifics (right). Mean with whiskers showing 95% confidence interval and distribution of C) Composite Z-score in Control-Alone, Control--Conspecifics, Experimental-Alone, and Experimental-Conspecifics. N = 10/ group. P value from unpaired Welch’s t-test.

To improve sensitivity of behavioral analysis and to allow comparison of the four groups of subjects, we used a Z-normalization procedure that integrates ^28^ these four parameters into a single score. A two-way analysis of variance of the Composite-Z score with conspecific presence and Schreckstoff exposure as between-subjects factors (Control-Alone, Control-Conspecific, Experimental-Alone and Experimental-Conspecific) showed a significant interaction (F (1, 36) = 23.79, p < 0.001). Follow-up Welch’s t-tests showed that the Composite-Z scores of subjects in Control-Alone and the Control-Conspecific groups were similar to each other (Figure 1C, p = 0.6). However, subjects in the Experimental-Alone group showed significantly higher Composite-Z scores than subjects in the Experimental-Conspecific group (Figure 1C, Supp. Figure 1, Supp. Video S1, S2; t(9.5) = 5.1, p < 0.001). This indicates that non-fearful conspecifics can socially buffer distressed subjects.

### Distressed subjects prefer familiar companions in a 2-choice assay

Familiarity with the conspecifics has recently been demonstrated as a factor that can influence effectiveness of social buffering in rodents ^12,29,30^. Hence, in the second and third experiments, we explored conspecific familiarity. Zebrafish are known to display a learned recognition of social partners in laboratory conditions ^31,32^. Building on this finding, we placed two groups of conspecifics in a two-compartment chamber behind the subject recovery tank (Figure 2A). Each group comprised of four non-fearful individuals. However, one group was familiar to the subject (raised in the same tank with the subject), while the other was unfamiliar (phenotypically distinguishable and raised in a different location). As in the first experiment, subjects were exposed to Schreckstoff, passed through a wash chamber, moved into an observation tank, and their behavior was recorded for 10 minutes. Subjects could see and swim freely between the two groups of conspecifics, but could not feel, hear or smell either. We quantified the time subjects spent in the two quadrants of the tank interfacing the two groups (Figure 2A). If distressed subjects had no preference, the time spent in these locations would not differ from that observable by chance (25% of total time). We found that subjects spent more time in the quadrant adjacent to familiar conspecifics than would be expected by chance (Figure 2B, mean = 40.0%, 95 CI [47.3%, 32.6%], t(15) = 3.66, p = 0.002). This was not the case for the time subjects spent in the quadrant adjacent to unfamiliar conspecifics (mean = 27.0%, 95 CI [32.3%, 21.6%, t(15) = 0.69, p = 0.5]). This indicates that subjects preferred familiar conspecifics during recovery. Additionally, it raises the possibility that familiarity facilitates social buffering in zerbafish.

**Figure 2.**
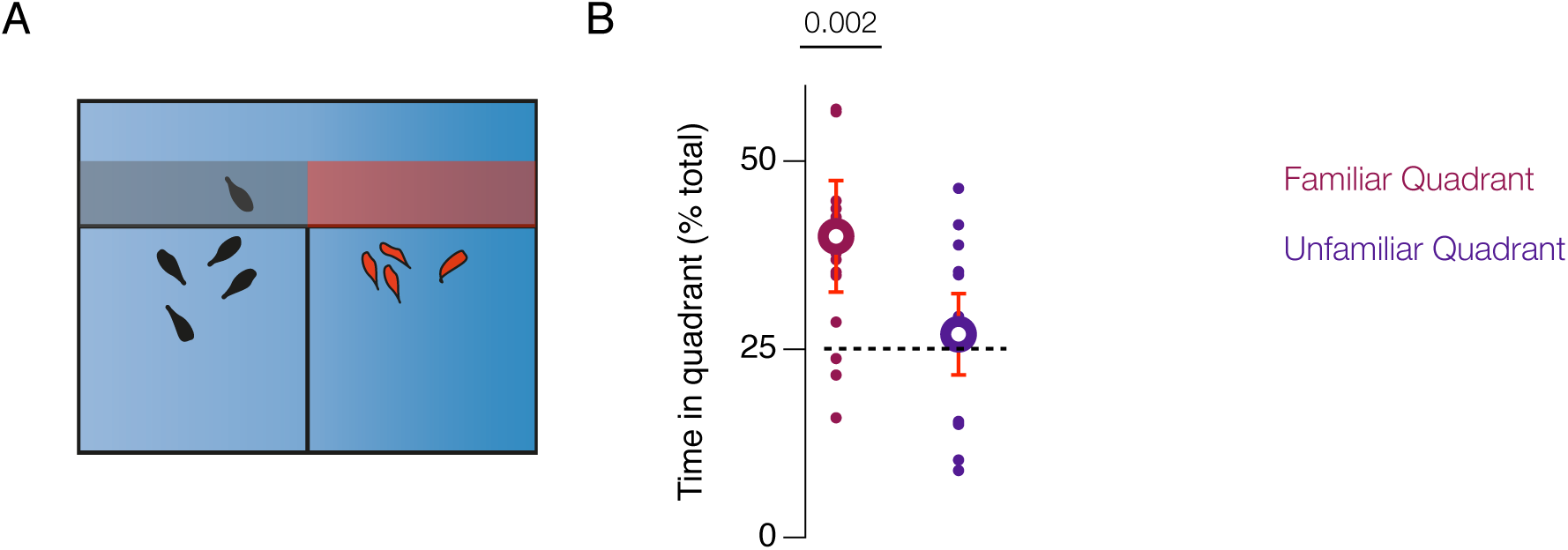
Distressed subjects prefer familiar companions in a 2-choice assay. A) Schematic showing the design of the social preference test. B) Mean with whiskers showing 95% confidence interval and distribution of subject’s time (percentage of total) in the quadrant adjacent to familiar (shaded gray in A) and unfamiliar conspecifics (shaded red in A). N = 16/group, p-values of unpaired Welch’s t-test comparing time in quadrants against chance (25% of total time, indicated by dashed line).

### Familiar conspecifics are effective in reducing plasma cortisol levels of distressed subjects

Stressors, such as threat of predation activate the hypothalamic-pitutary-adrenal axis (HPA axis) and the hypothalamic-pituitary-interrenal glands (HPI axis) in mammals and fish, respectively, thereby increasing circulating levels of plasma cortisol ^33,34^. Exposure to alarm substance is a stressor that increases circulating plasma cortisol levels in many species of fish including zebrafish ^27,35,36^. To test whether familiar conspecifics effectively dampen this response, subjects were exposed to the alarm substance and divided into three groups – those that recovered alone (Experiemental-Alone), those that recovered with familiar conspecifics (Experimental-Familiar), and those that recovered with unfamiliar conspecifics (Experimental-Unfamiliar). The Composite-Z score derived from the behavioral measures mentioned above was subjected to an ANOVA with group as a factor. A significant effect of group (F (2, 69) = 18.76, p < 0.001) was explored using unpaired Welch’s t-tests with modified Bonferroni correction for multiple comparisons. The tests revealed that subjects in the Experimental-Alone group (Figure 3A, mean = 0.8, 95 CI [1.2, 0.4]) differed significantly from both Experimental-Familiar and Experimental-Unfamiliar groups (Figure 3B, t(25.2, 32.3) = 5.1, 3.9, ps < 0.001), but the Experimental-Familiar (mean = −0.1, 95CI [0.2, −0.4]) and Experimental-Unfamiliar groups (mean = −0.1, 95 CI [0.1, −0.3]) did not differ from each other (t(33.1) = 1.6, p = 0.1). Therefore, both familiar and unfamiliar conspecifics promoted behavioral recovery from stress.

**Figure 3.**
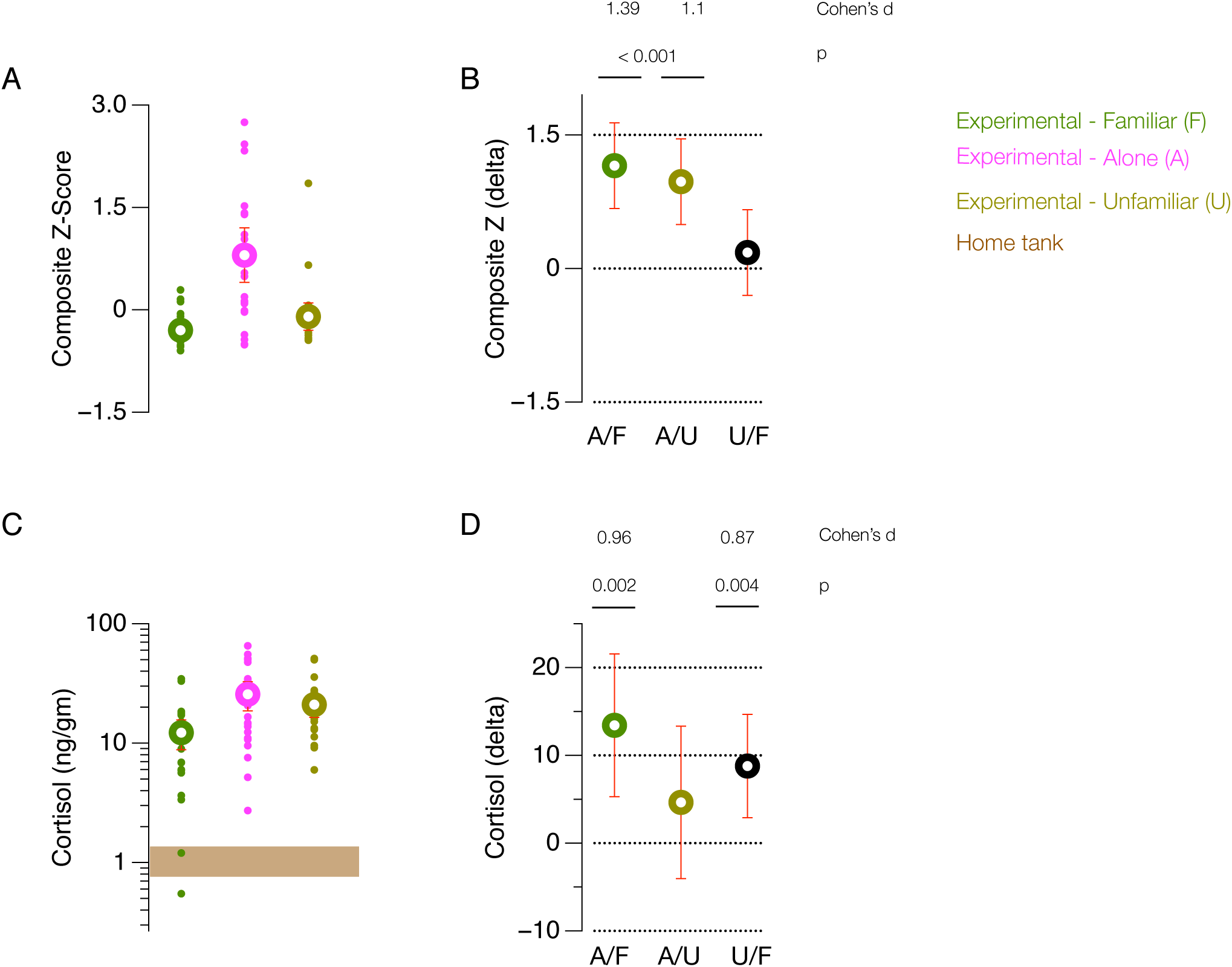
Familiar conspecifics reduce behavioral and physiological markers of distress. Mean with whiskers showing 95% confidence interval and distribution of A) Composite Z-scores and C) Whole-body cortisol levels in subjects from Experimental-Alone, Experimental-Familiar, and Experimental-Unfamiliar groups (N = 24/ group). Cortisol levels are represented in ng/gm of tissue. Difference of means (delta) represented as mean with whiskers showing 95% confidence interval for B) Composite Z-scores shown in A and D) Whole-body cortisol levels shown in C. Brown bar in C) shows the 95% confidence interval of cortisol level for fish collected from the home tank (N = 10). Effect size is computed as Cohen’s d and p values are adjusted for multiple comparisons. A/F: Experimental-Alone vs. Experimental-Familiar, A/U: Experimental-Alone vs. Experimental-Unfamiliar, U/F: Experimental-Unfamiliar vs. Experimental-Familiar.

An ANOVA of cortisol levels with group as a factor was also significant (F (2, 69) = 6.49, p =0.002). Follow-up unpaired Welch’s t-tests with modified Bonferroni correction for multiple comparison indicated that, compared to Experimental-Alone subjects (Figure 3C, mean = 25.6 ng/gm, 95 CI [32.7,18.6]), Experimental-Familiar subjects (Figure 3C, mean = 12.2 ng/gm, 95 CI [15.6,8.8]) showed a significant reduction in circulating plasma cortisol (Figure 3D, t(33.2) = 3.3, p = 0.002). Cortisol of Experimental-Familiar subjects was also lower than that of Experimental-Unfamiliar subjects (Figure 3D, t(42.5) = 3.0, p = 0.004). Experimental-Alone and Experimental-Unfamiliar subjects, on the other hand, did not differ (t(39.5) = 1.1, p = 0.28). Thus, familiarity of companions resulted in a faster suppression of stress induced HPI axis activation, further supporting similarities in the social buffering of zebrafish and some mammalian species.

### Central endogenous isotocin levels spike in the presence of familiar companions

Both stressors ^37^ and social interactions ^38^ can increase oxytocin release in the brain. Depending on the context, higher release is thought to facilitate greater social sensitivity ^39^ and/or counter HPA axis activation during stress ^8^. Exogenous administration of oxytocin or an oxytocin receptor antagonist has been shown to modulate HPA axis activity in humans, voles, and rats (reviewed in ^7^) and to regulate social interactions in zebrafish ^40^. Finally, a recent study demonstrated that social buffering by a pair-bonded male partner increased oxytocin release in the hypothalamus of stressed female prairie voles ^12^. These results prompted us to determine whether endogenous isotocin (zebrafish homologue of oxytocin) levels are affected in our assay. Thus, we measured isotocin from the heads of subjects that previously completed the Experimental-Alone, the Experimental-Familiar, and the Experimental-Unfamiliar conditions. Isotocin, like oxytocin is expected to have a short half-life of 6 to 9 minutes after release ^41^ and our measurements were terminal. We, hence, measured endogenous isotocin levels from two sets of subjects at two time-points (at 5, and at 10 minutes after subjects were moved into the observation tank).

An ANOVA with group as a factor was significant at both 5 minutes (F(2,45) = 6.3, p = 0.003) and 10 minutes (F(2, 40) = 5.1, p = 0.01). Follow-up unpaired Welch’s t-tests with modified Bonferroni correction for multiple comparisons revealed that at 5 minutes, isotocin levels were higher in Experimental-Familiar subjects (Figure 4A, B; mean = 22. 2 ng/fish, 95 CI [32.9, 11.5]) compared to Experimental-Alone (Figure 4B; mean = 6.8 ng/fish, 95 CI [8.9, 4.7], t(16.1) = 2.7, p = 0.014) and Experimental-Unfamiliar subjects (Figure 4B; mean = 8.2 ng/fish, 95 CI [11.6, 4.7], t(18.1) = 2.4, p = 0.025). Experimental-Alone and Experimental-Unfamiliar subjects were similar to each other (p = 0.52). At 10 minutes, isotocin levels were lower for the Experimental-Familiar group subjects (Figure 4C, D; mean = 2.2 ng/fish, 95 CI [2.7, 1.8]) relative to both Experimental-Alone (Figure 4D; mean = 6.1 ng/fish, 95 CI [8.8, 3.4], t(14.6) = 2.7, p = 0.017) and the Experimental-Unfamiliar subjects (Figure 4D; mean = 3.3 ng/fish, 95 CI [3.9, 2.6], t(21.9) = 2.5, p = 0.017).

**Figure 4.**
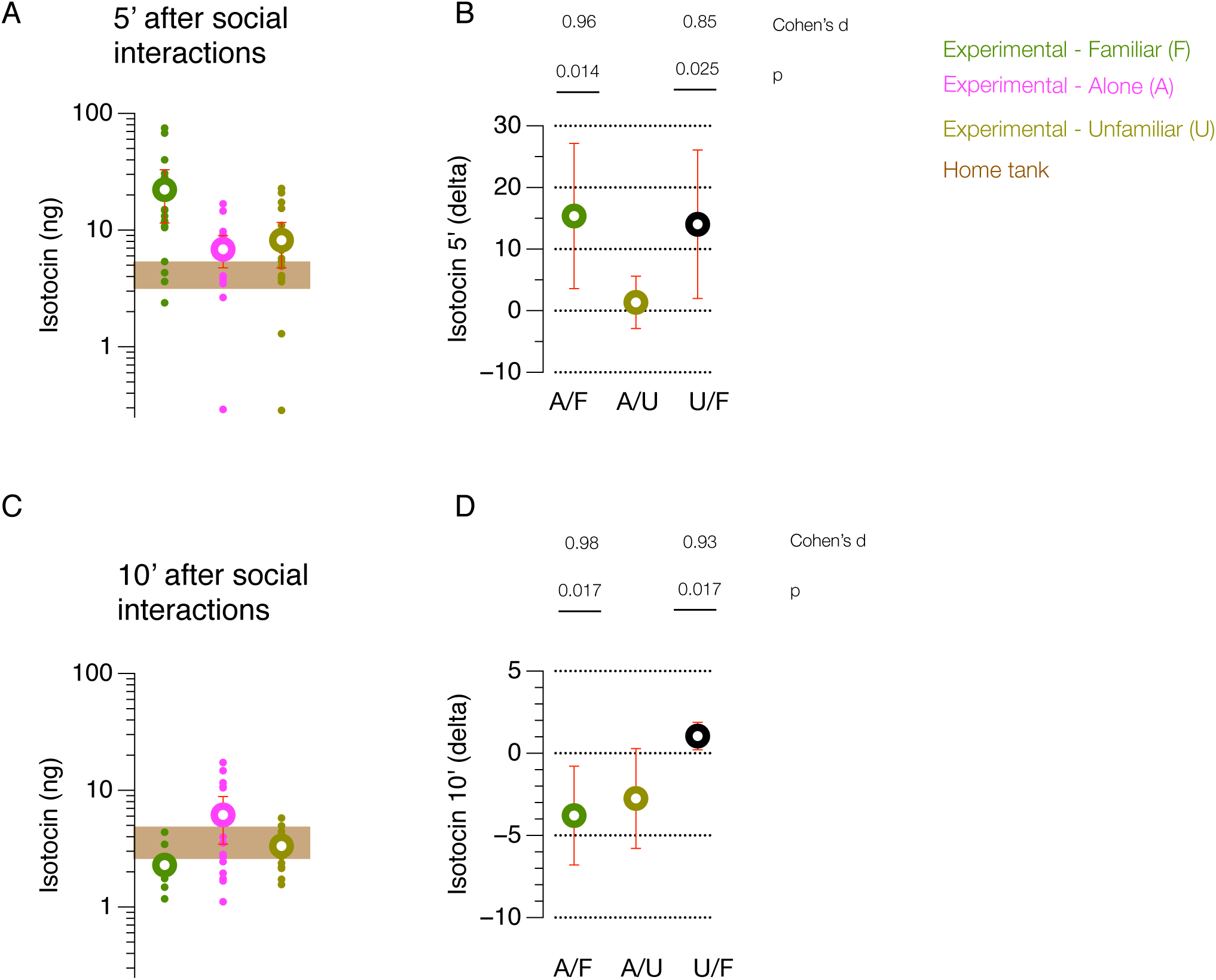
Endogenous isotocin levels spike after interactions with familiar companions. Mean with whiskers showing 95% confidence interval and distribution of isotocin levels (ng/fish) in subjects at A) 5 minutes (N = 16/group) and C) and at 10 minutes (N = 14/group) for groups Experimental-Alone, Experimental-Familiar, and Experimental-Unfamiliar. Brown bar shows the 95% confidence interval of isotocin level for fish collected from the home tank (N = 16). Difference of means (delta) represented as mean with whiskers showing 95% confidence interval in B) for Isotocin levels at 5 minutes shown in A and in D) for Isotocin levels at 10 minutes shown in C. Effect size is computed as Cohen’s d and p values are adjusted for multiple comparisons. A/F: Experimental-Alone vs. Experimental-Familiar, A/U: Experimental-Alone vs. Experimental-Unfamiliar, U/F: Experimental-Unfamiliar vs. Experimental-Familiar.

As isotocin levels in the Experimental-Familiar group were high at 5 minutes, we checked whether HPI axis suppression in Experiment 3 (Figure 3C, D) was also evident at this earlier time point (Supplementary Figure 2). Cortisol levels in all three groups of subjects however, were high and indistinguishable at 5 minutes. Together, these results suggest that an increase in central isotocin precedes HPI axis suppression when distressed subjects recover with familiar conspecifics, as compared to when recovering alone or with unfamiliar conspecifics.

## Discussion

We have shown that recovery from fear in zebrafish is accelerated by the presence of conspecifics, especially if they are familiar. These results provide important insights into the sociability of fish and point to the evolutionary continuity for basic social processes. Similar to the observations of effective buffering by a familiar conspecific in rats ^29^, zebrafish not only show behavioral recovery, but also display a faster suppression of HPI axis activity in the presence of familiar shoalmates. Further, familiar shoalmates increase central isotocin levels in subjects, which mirrors observations of increased oxytocin release in the presence of a bonded partner in the prairie vole ^12^. Zebrafish, like some other fish have been documented to recognize conspecific shoal members using visual cues ^31,32^. Experiments here suggest that this preference is unaltered even when they are fearful. This in turn suggests that like many mammalian species, zebrafish benefit from established social connections when facing danger. Although these benefits and the related representations of self and other are undoubtedly much more simple in gregarious fish than they are in mammals, they nevertheless exist and, as shown here, play a role in fish emotion and behavior. Among the complex set of determinants including learned preferences ^31,32^ and active decision-making ^42^ that influence group membership in fish shoals ^43^, the stress relief that comes from familiarity may be an additional factor.

In our hands, the presence of unfamiliar conspecifics resulted in behavioral recovery without suppression of the HPI axis. This is consistent with an observation in rats of a decrease in some behavioral signs of stress in the presence of an unfamiliar conspecific ^29^. It is also consistent with an observation that glucocorticoid receptor mutant zebrafish show behavioral recovery from immobility without a decrease in blood-level cortisol when presented with a conspecific visual ^19^. These findings imply that behavioral measures may not correlate accurately with physiological measures of distress. One possibility is that the behavior of the subject in these situations reflects the “selfish-herd” phenomenon. Subjects may simply act to blend in with a group to avoid predation when threatened ^44^. Alternatively, it is also possible that currently used behavioral parameters represent the state of distress incompletely. Additional parameters such as a subtle change in body posture, if quantifiable, might reveal differences in the subjects’ behavior that are otherwise obscured.

Social buffering, as used in mammals implies active participation or action of the companions that goes beyond being present (such as grooming and vocalizing). In the experiments reported here, we focused on the subject behavior and did not analyze the behavior of companions in detail. Whether companions behave differently when they encounter a fearful individual, or actively contribute to the recovery of the subject was not studied. Further experiments are required to evaluate if the behavior of companions, as a group or as individuals within a group changes, to influence the recovery of the subject. This will add to the growing body of research focusing on the social cognitive abilities of fishes (reviewed in ^45-47^).

Behaviors that require both social and individual recognition such as transitive inference and goal-directed cooperation have already been documented in fishes ^48,49^. Moreover, an evolutionarily conserved set of brain nuclei that form a social-decision making network and are common to all vertebrate brains have also been identified ^50^. Our results advance this literature ^51^ by revealing a previously unanticipated complexity in the social processes of fish. Overall, the high degree of similarity in these processes between mammals and fish points to a likely common ancestry. This has important theoretical and practical implications, but requires further investigations. Specifically, it has the potential to address the current interest in understanding the mechanisms that produce and sustain social bonds. Are there fundamental processes that evolved once in the vertebrate lineage and subsequently fortified by species-specific mechanisms? In spite of the recognition that this is a priority research area due to the influence social ties have on human health ^52^, insights into the genetic, molecular and neural basis of attachments have proven difficult because ethical, technological, and financial constraints limit research possibilities. A non-mammalian model organism with a capacity for mammalian-like social behavior offers new avenues, and promises discoveries rendered impossible by research with an exclusively mammalian focus.

## Materials and Methods

### Fish lines

All experiments were conducted in accordance with guidelines approved by Institutional Animal Care and Use Committee of Biopolis (#100594). Subjects were adult fish (4 to 8 months old) in AB background, either heterozygous or homozygous for *nacre*. At present it is not clear if the pigment mutation *nacre* may have any role in the behavior described, but subject fish were pseudo randomly assigned to different conditions to get approximately equal number of heterozygotes and homozygotes. Initial experiments suggested potential sex-dependent differences in recovery rate, however, these differences did not meet the criteria for significance. Therefore, equal number of males and females were used in each experiment. To facilitate tracking, subjects were raised with conspecifics with different skin pigmentation and were randomly assigned to one of the conditions in an experiment. To allow the experimenter to distinguish subjects from companions, conspecific companions included Danio Reds (expressing a red fluorescent protein in the muscle tissue ^53^) or Wku – a spontaneous pigment mutation isolated in the fish facility. For experiments requiring familiar conspecifics, subjects and conspecifics were raised together from the time of hatching. Unfamiliar conspecifics were raised in a separate aquarium rack, such that they were never seen or encountered by subjects before the experiment. Please note that the empirical data shown here was collected in the context of a larger project. Behavioral and cortisol data from 11 of the 24 subjects in the Experimental-Alone condition in the third experiment (Figure 3) was previously reported and published here ^27^.

### Behavioral assays

Conspecifics (2 males and 2 females) were collected from the facility and transferred into a custom-made glass tank (30 cm x 6 cm x 13 cm -L x W x H) filled with aquarium water to a depth of 10 cm. The tank was uniformly illuminated from above using a white light LED (i-bar LED lamp, Koncept) in a darkened room, such that the experimenter was obscured. Conspecific fish were acclimatized for 8‘-10’. Subject fish were then collected from the facility and placed in a beaker with 50ml aquarium water. 200 ul of 1X Schreckstoff (see below) was introduced into this beaker for the Experimental subjects. An equivalent volume of aquarium water was used for Control subjects. After 2’, subjects were transferred to a wash chamber with 400 ml aquarium water via a net so that alarm substance containing water was discarded. Subjects were then transferred into a 100 ml beaker with 50 ml aquarium water via a net such that wash chamber water was also discarded. They were then gently delivered into the observation chamber. Subject behavior was recorded for 10’ on a MacBook with Agent v5 HD web camera placed ~50 cm in front of the tank

For social choice experiments, conspecifics were placed in a two-compartment chamber (each compartment was10 cm x 15 cm x 13 cm – L x W x H) behind the subject tank. Unfamiliar and familiar conspecifics could see and interact with the subject but not with each other. Tanks were illuminated from the bottom with white light using a lightbox (LED GraphPad). Videos were recorded on Flea 3 (Point Grey) placed ~75 cm above the tanks using a 16 mm Computar lens (COM1614MP2).

In experiments that required cortisol or isotocin measurement, behavioral experiments were always performed between 14:00–18:00 in the afternoon. Subject fish were netted at the end of the observation period, immediately euthanized in ice water, dried using Kimwipes, and flash frozen in dry ice. Samples were kept at −70°C until thawed for cortisol or isotocin extraction.

### Preparation of 1X Alarm Pheromone (Schreckstoff)

Schreckstoff was prepared as described in ^54^ with some modifications. Briefly, 4 to 5 fish were euthanized by immersion in ice-cold water. 7 to 10 shallow lesions were made on the skin of each euthanized zebrafish, using a sharp knife (Sharpoint, 22.5° stab), such that drawing of blood was minimal. Fish were then immersed in 2 ml of 20 mM Tris-Cl (pH 8.0) and rocked on a rocker for 5’. The 2 ml crude extract was heated at 95°C overnight, then centrifuged at 13.2k rpm in an Eppendorf microfuge for 10’. The supernatant and used as 1X Schreckstoff.

### Analysis

Videos were recorded at 25 fps and re-digitized at 12 fps. Tracking of the subject fish position through the video was automated using the “track objects” algorithm in MetaMorph. 6.3. Macros in Microsoft Excel were written to derive the position of the fish in the arena (swimmable tank area) and speed (in mm/sec) followed by automatic computation of the following dependent variables - percentage time spent in the bottom quarter area of the tank, pausing, freezing and darting episodes as described ^27^. Briefly, pausing episodes were defined as 1 second of immobility (speed<3.5 mm/sec). Freezing episodes were defined as extended periods of pausing lasting at least 5” or more. Darting episodes were defined as erratic swimming episodes where the swim speed exceeded the normal swimming speed of a fish by 10 SD. This normal swimming speed was defined as the average swimming speed of 5 fish (3 males and 2 females) observed in the same tank for 10 minutes each. In the social preference experiment (Experiment 2), the area of the subject tank was divided into quadrants and total time in each quadrant was quantified.

To compute the Composite Z-score, we first normalized data across all conditions for each dependent variable (DV; Supplementary Figure S1) by calculating z-scores for each observation. That is, we computed z = (X - μ)/σ, where X = observation, μ = population mean, and σ = population standard deviation. For normalization one of the strategies as recommended by ^55^ was used. Since subjects in the Control conditions rarely displayed certain DVs (such as pausing, darting), population μ and σ for a DV included Experimental and Control conditions and Controls therefore show negative scores. We then used an integrated behavioral z-score system as recommended in ^28^, that is composite z score = (z_DV1_ + z _DV2_ + z _DV3_ + z_DV4_) / 4 (# of DVs).

### Cortisol extraction and assay

Whole-body cortisol was prepared as described before with some modifications ^56^. Briefly, fish were thawed from −70°C, weighed and decapitated. The fish body without the head was dissected into five pieces and divided into five 2ml Eppendorf tubes. 200ul of 1X PBS (pH 7.2) at 4°C was added to each tube and homogenized using an Ultra-Turrax Disperser (T10 basic; IKA). Cortisol was extracted in 1400 ul of Ethyl Acetate (Sigma) by vortexing the tubes for 30”, followed by centrifugation at 7000g for 15’ at room temperature. Organic layer (top) that contained cortisol was collected in fresh tubes and left in the fume hood overnight to allow ethyl acetate to evaporate. Cortisol was reconstituted in 1ml of 1X PBS and stored at 4°C. ELISA was performed using either Cayman (Item # 500360) or Enzo (Catalog # ADI-900-071) cortisol measurement kits following the manufacturer’s instructions within 2-3 days of extraction. Cortisol levels of samples were determined by constructing a standard curve and interpolation.

### Isotocin extraction and assay

Fish were thawed from −70°C, dried, decapitated and the head was weighed. Peptide extraction was performed from the head by reverse phase chromatography using disposable Sep-Pak C_18_ cartridges (Millipore) as described ^57^ with some modifications. Briefly, heads were homogenized using an Ultra-Turrax Disperser (T10 basic; IKA) in 1ml of 0.25% Acetic acid (v/v) for 30”. The homogenate was placed in boiling water bath for 200”, cooled on ice and centrifuged at 1500 RPM for 30’ at 4°C. Sep-Pak C_18_ cartridge was activated by pushing 5ml of 100% Methanol followed by 20ml of distilled water. Supernatant from tissue homogenate was pushed through the column slowly over 2 minutes. The column was then washed by pushing 10 ml of 4% Acetic Acid (v/v) followed by 10 ml of distilled water. Neuropeptides were then eluted by passing 1 ml of 3mM HCl in 100% Ethanol through the column over 2 minutes. The eluate was flash frozen in dry ice and evaporated to dryness using a centrifugal concentrator under vacuum and stored at −20°C till use. Determination of isotocin was performed using Enzo ELISA kit (Enzo, NY, USA) following manufacturer’s instructions within 10 days of extraction. Isotocin levels of samples were determined by constructing a standard curve and interpolation. Intra-assay variance was calculated to be between 1% −7.5%. Isotocin detection of 2 subjects each in the Experimental-Familiar, Experimental-Unfamiliar groups and 1 subject from the Experiment-Alone at the 10-minute time point failed and could not be included.

### Statistics

For all experiments distribution of individual data points, mean and the error bars showing 95% confidence interval (95 CI) are presented as recommended ^58^. Experimental data was analyzed by one-way or two-way analysis of variance followed by two-sided unpaired Welch’s t-tests. Differences were considered significant when p < 0.05. In experiments 3 and 4, modified Bonferroni correction was applied as suggested in ^59^ for multiple comparisons to set the alpha threshold of p value to 0.033.

## Acknowledgments

We thank J Huang and X Y Koh for assistance with behavioral experiments. We also thank Seetha Krishnan for assistance with analysis of behavior data. This work was funded by the Biomedical Research Council, Singapore.

## Author contributions

Project was conceived by ASM, AS and SJ. ASM and NPYT performed experiments. ASM and CK performed cortisol and isotocin extractions. ASM and RKC formulated and wrote codes for automated data analysis. AS and ASM designed experiments, analyzed the data and wrote the manuscript with assistance from SJ.

## Supplementary Figure Legends

**Supp. Figure 1.**
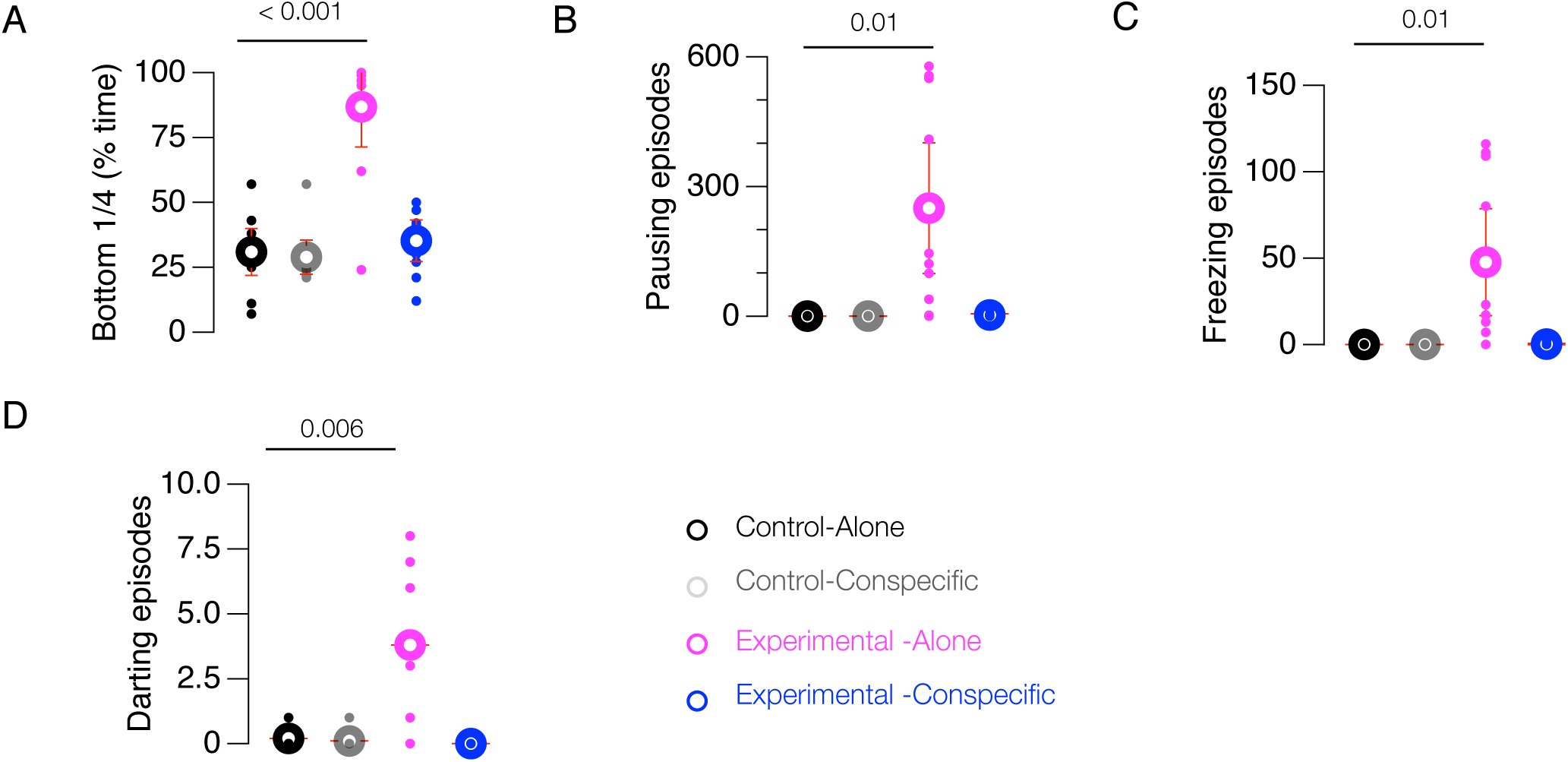
Dependent variables integrated to derive composite Z-score in Figure 1. Dependent variables showing A) percent time in the bottom 1/4^th^ of the tank, B) pausing episodes, C) freezing episodes and D) darting episodes in Control – alone, Control-with conspecifics, Experimental-Alone, and Experimental - with conspecifics. P-values are calculated from Unpaired T-tests.

**Supp. Figure 2.**
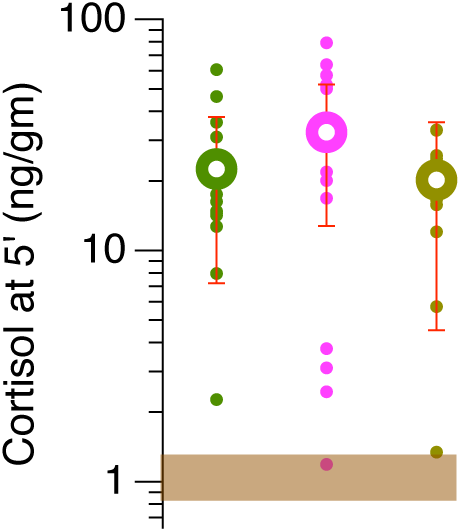
Cortisol levels are high and similar between groups at 5 minutes. Whole-body cortisol levels in subjects from Experimental-Alone, Experimental-Familiar and Experimental-Unfamiliar groups (N = 16/group). Brown bar shows the 95% confidence interval of cortisol level for fish collected from the home tank.

## REFERENCES

1. Alexander, R. D. The evolution of social behavior. Annual review of ecology and systematics 5, 325–383 (1974).

2. Rubenstein, D. I. On predation, competition, and the advantages of group living. Perspectives in ethology 3, 205–231 (1978).

3. Krause, J. & Ruxton, G. D. Living in groups. (Oxford University Press, USA, 2002).

4. Sapolsky, R. M. The Influence of Social Hierarchy on Primate Health. Science (New York, NY) 308, 648–652 (2005).

5. Davitz, J. R. & Mason, D. J. Socially facilitated reduction of a fear response in rats. J Comp Physiol Psychol 48, 149–151 (1955).

6. Kikusui, T., Winslow, J. T. & Mori, Y. Social buffering: relief from stress and anxiety. Philosophical Transactions of the Royal Society B: Biological Sciences 361, 2215–2228 (2006).

7. Hennessy, M. B., Kaiser, S. & Sachser, N. Social buffering of the stress response: diversity, mechanisms, and functions. Frontiers in Neuroendocrinology 30, 470–482 (2009).

8. Smith, A. S. & Wang, Z. Salubrious effects of oxytocin on social stress-induced deficits. Hormones and Behavior 61, 320–330 (2012).

9. Panksepp, J., Herman, B. H., Vilberg, T., Bishop, P. & DeEskinazi, F. G. Endogenous opioids and social behavior. Neuroscience & Biobehavioral Reviews 4, 473–487 (1980).

10. Carter, C. S. Neuroendocrine perspectives on social attachment and love. Psychoneuroendocrinology 23, 779–818 (1998).

11. Neumann, I. D. Involvement of the brain oxytocin system in stress coping: interactions with the hypothalamo-pituitary-adrenal axis. Prog. Brain Res. 139, 147–162 (2002).

12. Smith, A. S. & Wang, Z. Hypothalamic oxytocin mediates social buffering of the stress response. Biological Psychiatry 76, 281–288 (2014).

13. Remage-Healey, L., Adkins-Regan, E. & Romero, L. M. Behavioral and adrenocortical responses to mate separation and reunion in the zebra finch. Hormones and Behavior 43, 108–114 (2003).

14. Hennessy, M. B., Hornschuh, G., Kaiser, S. & Sachser, N. Cortisol responses and social buffering: a study throughout the life span. Hormones and Behavior 49, 383–390 (2006).

15. Coe, C. L., Mendoza, S. P., Smotherman, W. P. & Levine, S. Mother-infant attachment in the squirrel monkey: adrenal response to separation. Behav Biol 22, 256–263 (1978).

16. Coe, C. L., Franklin, D., Smith, E. R. & Levine, S. Hormonal responses accompanying fear and agitation in the squirrel monkey. Physiol Behav 29, 1051–1057 (1982).

17. Kirschbaum, C., Klauer, T., Filipp, S. H. & Hellhammer, D. H. Sex-specific effects of social support on cortisol and subjective responses to acute psychological stress. Psychosom Med 57, 23–31 (1995).

18. Hennessy, M. B., O’Leary, S. K., Hawke, J. L. & Wilson, S. E. Social influences on cortisol and behavioral responses of preweaning, periadolescent, and adult guinea pigs. Physiol Behav 76, 305–314 (2002).

19. Ziv, L. et al. An affective disorder in zebrafish with mutation of the glucocorticoid receptor. Mol. Psychiatry (2012). doi:10.1038/mp.2012.64

20. Allen, P. J., Barth, C. C., Peake, S. J., Abrahams, M. V. & Anderson, W. G. Cohesive social behaviour shortens the stress response: the effects of conspecifics on the stress response in lake sturgeon Acipenser fulvescens. Journal of Fish Biology 74, 90–104 (2009).

21. Hamilton, W. D. Geometry for the selfish herd. J. Theor. Biol. 31, 295–311(1971).

22. Faustino, A. Social buffering of fear in zebrafish. (2016).

23. Spence, R., Ashton, R. & Smith, C. Oviposition decisions are mediated by spawning site quality in wild and domesticated zebrafish, Danio rerio. Behaviour 144, 953–966 (2007).

24. Neumann, I. D. Brain oxytocin: a key regulator of emotional and social behaviours in both females and males. J. Neuroendocrinol. 20, 858–865 (2008).

25. Graustella, A. J. & MacLeod, C. A critical review of the influence of oxytocin nasal spray on social cognition in humans: evidence and future directions. Hormones and Behavior 61, 410–418 (2012).

26. Frisch, von, K. Zur psychologie des fisch-schwarmes. Naturwissenschaften 26, 601–606 (1938).

27. Schirmer, A., Jesuthasan, S. & Mathuru, A. S. Tactile stimulation reduces fear in fish. Frontiers in behavioral neuroscience 7, 167 (2013).

28. Guilloux, J.-P., Seney, M., Edgar, N. & Sibille, E. Integrated behavioral z-scoring increases the sensitivity and reliability of behavioral phenotyping in mice: relevance to emotionality and sex. J Neurosci Methods 197, 21–31 (2011).

29. Kiyokawa, Y., Honda, A., Takeuchi, Y. & Mori, Y. A familiar conspecific is more effective than an unfamiliar conspecific for social buffering of conditioned fear responses in male rats. Behavioural Brain Research 267, 189–193 (2014).

30. Bowen, M. T. et al. Defensive aggregation (huddling) in Rattus norvegicus toward predator odor: individual differences, social buffering effects and neural correlates. PLoS ONE 8, e68483 (2013).

31. Engeszer, R., Ryan, M. & Parichy, D. Learned social preference in zebrafish. Current Biology 14, 881–884 (2004).

32. Griffiths, S. W. & Ward, A. Learned recognition of conspecifics. Fish Cognition and Behavior (2007).

33. Szabo, S., Tache, Y. & Somogyi, A. The legacy of Hans Selye and the origins of stress research: a retrospective 75 years after his landmark brief ‘letter’ to the editor# of nature. Stress 15, 472–478 (2012).

34. Mommsen, T., Vijayan, M. & Moon, T. Cortisol in teleosts: dynamics, mechanisms of action, and metabolic regulation. Reviews in Fish Biology and Fisheries 9, 211–268 (1999).

35. Sanches, F. H. C., Miyai, C. A., Pinho-Neto, C. F. & Barreto, R. E. Stress responses to chemical alarm cues in Nile tilapia. Physiol Behav 149, 8–13 (2015).

36. Rehnberg, B. G. & Schreck, C. B. Chemosensory detection of predators by coho salmon (Oncorhynchus kisutch): behavioural reaction and the physiological stress response. Canadian Journal of Zoology 65, 481–485 (1987).

37. Carter, C. S., Grippo, A. J., Pournajafi-Nazarloo, H., Ruscio, M. G. & Porges, S. W. Oxytocin, vasopressin and sociality. Prog. Brain Res. 170, 331–336 (2008).

38. Waldherr, M. & Neumann, I. D. Centrally released oxytocin mediates mating-induced anxiolysis in male rats. Proc Natl Acad Sci USA 104, 16681–16684 (2007).

39. Carter, C. S. Oxytocin pathways and the evolution of human behavior. Annual review of psychology 65, 17–39 (2014).

40. Braida, D. et al. Neurohypophyseal hormones manipulation modulate social and anxiety-related behavior in zebrafish. Psychopharmacology 220, 319–330 (2012).

41. Churchland, P. S. & Winkielman, P. Modulating social behavior with oxytocin: how does it work? What does it mean? Hormones and Behavior 61, 392–399 (2012).

42. Engeszer, R., Da Barbiano, L., Ryan, M. & Parichy, D. Timing and plasticity of shoaling behaviour in the zebrafish, Danio rerio. Animal Behaviour 74, 1269–1275 (2007).

43. Parrish, J. K. & Edelstein-Keshet, L. Complexity, pattern, and evolutionary trade-offs in animal aggregation. Science (New York, NY) 284, 99–101 (1999).

44. King, A., Wilson, A., Wilshin, S. & Lowe, J. Selfish-herd behaviour of sheep under threat. Current Biology (2012).

45. Oliveira, R. F. Mind the fish: zebrafish as a model in cognitive social neuroscience. Front Neural Circuits 7, 131 (2013).

46. Bshary, R. & Brown, C. Fish cognition. Curr Biol 24, R947–50 (2014).

47. Bshary, R., Gingins, S. & Vail, A. L. Social cognition in fishes. Trends in Cognitive Sciences 18, 465–471 (2014).

48. Grosenick, L., Clement, T. S. & Fernald, R. D. Fish can infer social rank by observation alone. Nature 445, 429–432 (2007).

49. Vail, A. L., Manica, A. & Bshary, R. Fish choose appropriately when and with whom to collaborate. Curr Biol 24, R791–3 (2014).

50. O’Connell, L. A. & Hofmann, H. A. Evolution of a vertebrate social decision-making network. Science (New York, NY) 336, 1154–1157 (2012).

51. Barrett, L., Henzi, P. & Rendall, D. Social brains, simple minds: does social complexity really require cognitive complexity? Philos Trans R Soc Lond B Biol Sci 362, 561–575 (2007).

52. Eisenberger, N. I. & Cole, S. W. Social neuroscience and health: neurophysiological mechanisms linking social ties with physical health. Nature Neuroscience 15, 669–674 (2012).

53. Wan, H. et al. Generation of two-color transgenic zebrafish using the green and red fluorescent protein reporter genes gfp and rfp. Mar. Biotechnol. 4, 146–154 (2002).

54. Mathuru, A. S. et al. Chondroitin Fragments Are Odorants that Trigger Fear Behavior in Fish. Curr Biol 22, 538–544 (2012).

55. Birmingham, A. et al. Statistical methods for analysis of high-throughput RNA interference screens. Nat Meth 6, 569–575 (2009).

56. De Marco, R. J., Groneberg, A. H., Yeh, C.-M., Castillo Ramírez, L. A. & Ryu, S. Optogenetic elevation of endogenous glucocorticoid level in larval zebrafish. Front Neural Circuits 7, 82 (2013).

57. Pierson, P. M., Guibbolini, M. E., Mayer-Gostan, N. & Lahlou, B. ELISA measurements of vasotocin and isotocin in plasma and pituitary of the rainbow trout: effect of salinity. Peptides 16, 859–865 (1995).

58. Krzywinski, M. & Altman, N. Points of Significance: Error bars. Nat Meth 10, 921–922 (2013).

59. Keppel, G. & Wickens, T. D. Design and Analysis: A Researcher’s Handbook. (Prentice Hall, 2004).

